# Misregulation of MYB16 causes stomatal cluster formation by disrupting polarity in asymmetric cell division

**DOI:** 10.1101/2021.05.03.442461

**Authors:** Shao-Li Yang, Ngan Tran, Meng-Ying Tsai, Chin-Min Kimmy Ho

## Abstract

Stomata and leaf cuticle regulate water evaporation from the plant body and balance the trade-off between photosynthesis and water loss. We identified MYB16, a key transcription factor controlling cutin biosynthesis, from previous stomatal lineage ground cell (SLGC)-enriched transcriptome study. The preferential localization of MYB16 in SLGCs but not meristemoids suggests a link between cutin synthesis and stomatal development. Here, we showed that downregulation of MYB16 in meristemoids was directly mediated by the stomatal master transcription factor, SPEECHLESS (SPCH). The suppression of MYB16 before asymmetric division was crucial for stomatal patterning because overexpression or ectopic expression of MYB16 in meristemoids increased impermeability and elevated stomatal density and clusters. The aberrant pattern of stomata was due to reduced and disrupted establishment of polarity during asymmetric cell division. Manipulating polarity by growing seedlings on hard agar rescued stomatal clusters and polarity defects in MYB16 ectopic lines. By expressing a cutinase in MYB16 ectopic lines, stomatal clustering was reduced, which suggests that the ectopic accumulation of cuticle affects the polarity in asymmetrically dividing cells and causes clustered stomata. Taken together, inhibiting MYB16 expression by SPCH in early stomatal lineage is required to correctly place the polarity complex for proper stomatal patterning during leaf morphogenesis.

## INTRODUCTION

Terrestrialization is a critical event by which organisms moved from the ocean to the land. In plants, two pivotal features — stoma and the cuticle layer — evolved to adapt to major changes from water to air. Stomata are valves on the plant surface to control gas exchange and water loss. The cuticle layer is a barrier between the external environment and air-exposed plant surface. The coordination of stomata and cuticle on the epidermis balances the trade-off between photosynthesis and water evaporation.

In *Arabidopsis*, stomata are created by a series of asymmetric and oriented cell divisions to ensure proper stomatal density and distribution on epidermis. Stomatal lineages are initiated from the expression of SPEECHLESS (SPCH), the basic helix-loop-helix transcription factor, to drive asymmetric cell division and produce meristemoids, precursors of guard cells (MacAlister et al., 2007), and stomatal lineage ground cells (SLGCs), which may become pavement cells or reinitiate asymmetric division to produce more stomata. The stomatal pattern follows a “one-cell-spacing” rule, which means that two stomata never directly contact each other (Geisler et al., 2000). Two pathways regulate stomatal patterning. One relies on cell–cell communication and the other is the establishment of polarity during asymmetric cell division. EPIDERMAL PATTERNING FACTOR (EPF) family peptide ligands including EPF1 and EPF2 are secreted from stomatal lineage cells to restrict stomatal development of neighbor cells (Hara et al., 2007; Hunt and Gray, 2009). Upon directly binding to the receptor-like kinase ERECTA family (ERf) (Lee et al., 2012), the EPF signal activates the mitogen-activated protein kinase (MAPK) cascade (Wang et al., 2007) to diminish SPCH stability (Lampard et al., 2008), which further prevents cells from entering the stomatal lineage. During asymmetric division, the polarity complex is formed in SLGCs and is integrated with MAPK signalling to decrease SPCH level, ultimately breaking the stomatal fate in SLGCs (Zhang et al., 2015). The complex includes BREAKING OF ASYMMETRY IN THE STOMATAL LINEAGE (BASL), POLAR LOCALIZATION DURING ASYMMETRIC DIVISION AND REDISTRIBUTION (POLAR) and the BREVIS RADIX (BRX) family. These factors interact with each other and form a crescent to recruit MAPK components in SLGCs (Dong et al., 2009; Houbaert et al., 2018; Pillitteri et al., 2011; Rowe et al., 2019; Zhang et al., 2015). Thus, the combination of cell–cell communication, MAPK signaling and the polarity complex regulates stomatal patterning in the epidermis.

As compared with the dynamic behavior of stomata, cuticle layers form a physical barrier to protect tissue against dehydration. A cuticle layer consists of different kinds of lipid polymers. Cutin and waxes are two major polymer components conserved across plant species (Yeats and Rose, 2013; Bhanot et al., 2021). In *Arabidopsis*, cutin monomers are synthesized in the endoplasmic reticulum (ER) membrane, transported to the outside of cells and polymerized to become a thin layer in epidermis. LONG-CHAIN ACYL-COA SYNTHETASE 1 (LACS1) and LACS2 are essential for transforming C16/C18 fatty acid into acyl-CoA precursors for both cutin and waxes (Lü et al., 2009). Following ω-hydroxylation and midchain hydroxylation by the cytochrome P450 enzymes CYP86A4 and CYP77A6 (Li-Beisson et al., 2009), acyl-CoA precursors are catalyzed by the GLYCEROL-3-PHOSPHATE SN-2-ACYLTRANSFERASE (GPAT) family to produce a cutin monomer, 2-monoacylglycerol (2-MAG) (Yang et al., 2010, 2012). After leaving the ER membrane, 2-MAG is then transported into the extracellular matrix (ECM) by the ATP-binding cassette (ABC) transporters ABCG11 and ABCG32 (McFarlane et al., 2010; Bessire et al., 2011). Finally, ECM-localized cutin synthase polymerizes the exported 2-MAG to form cuticle layers.

Although most cutin biosynthesis-related genes are identified or predicted as biosynthesis enzymes, few transcription factors have been found to activate the pathway. *WAX INDUCER1/SHINE1* (*WIN1/SHN1*), an AP2 domain-containing transcription factor, was first identified from an overexpression screen and found to control wax formation (Aharoni et al., 2004; Broun et al., 2004). MYB16, an R2R3 MYB transcription factor, together with MYB106 from the same MYB subgroup (Stracke et al., 2001), could promote petal cell outgrowth and directly upregulate the expression of genes involved in cutin biosynthesis (Baumann et al., 2007; Oshima et al., 2013), which suggests a link between cuticle development and epidermal cell differentiation. In barley, the wax-deficient *eceriferum-g* mutant *cer-g* features clustered stomata (Zeiger and Stebbins, 1972). A mutation in *HIC (HIGH CARBON DIOXIDE),* an enzyme involved in the synthesis of very-long chain fatty acid, produced a thinner cuticle and higher stomatal density in epidermis (Gray et al., 2000). Also, overexpression of *WIN1/SHN1* decreased stomatal density (Yang et al., 2011). This evidence suggests a connection between stomatal pattern and cuticle defects.

Cutin may carry inhibitors or directly implement an inhibitory signal for stomatal development (Bird and Gray, 2003). Also, a cuticle layer may provide an elastic shell to regulate mechanical properties and further affect plant development (Galletti et al., 2016). Cell division and cell expansion during growth deforms the cell membrane and extracellular layer to generate surface mechanical force. Disruption of ECM reduces cell adhesion and results in disorganized tumor-like growth on shoot (Krupková et al., 2007; San-Bento et al., 2014); therefore, the tissue-wide coordination of cell–cell adhesion is important to maintain the proper patterning of a tissue. Moreover, the localization of polarized proteins could also respond to the action force from the growth direction in both leaf and shoot (Heisler et al., 2010; Bringmann and Bergmann, 2017). Altered mechanical force in leaf epidermis by laser ablation leads to the redistribution of polarity proteins (Bringmann and Bergmann, 2017). In animals, elevated mechanical stress in neutrophil cells led to failed polar component recruitment and finally cessation of cell migration (Houk et al., 2012). Plants have no well-established method to manipulate the mechanical force without damaging cells, but growing seedlings on high-percentage agar plates reduced tensile stress on epidermis and rescued cell–cell gaps between epidermal cells in a cell adhesion mutant, *quasimodo1* (*quo1*) (Verger et al., 2018). Cuticle is one type of ECM. Despite no direct evidence linking mechanical force and cuticular layer on epidermis, observations from tomato fruit showed a positive correlation between cuticle thickness and stiffness (Matas et al., 2004), so the amount of cuticle affects mechanical properties on cells.

Transcriptome profiling of stomatal lineage cells and expression analysis revealed enrichment and preferential localization of *MYB16* in SLGCs, which raises the question of why MYB16 prefers SLGCs but not meristemoids (Ho et al., 2021). To understand the role of MYB16 in stomatal formation, here we investigated the dynamics of SPCH and MYB16 expression during asymmetric cell division and found that *MYB16* was transcriptionally downregulated by SPCH in meristemoids. Overexpression or ectopic expression of MYB16 in an early stomatal lineage resulted in clustered stomata and increased amount of cuticle in epidermis. The disrupted stomatal pattern was due to the decreased and mis-polarized polarity protein during asymmetric division. Suboptimal water potential conditions or ectopically expressing a cutinase gene, CUTICLE DESTRUCTING FACTOR 1 (CDEF1) (Takahashi et al., 2010), partially rescued stomatal clusters in the MYB16 ectopic expression line. This finding suggests that the accumulation of cutin modulates mechanical properties, thereby affecting the polarity protein behavior. Appropriate MYB16 regulation in asymmetrically dividing cells is required to establish proper polarity for accurate stomatal patterning.

## RESULTS

### Preferential localization of MYB16 in SLGCs is due to negative regulation by SPCH in meristemoids

Previous study has shown that *MYB16* transcripts are enriched in SLGCs and MYB16 protein is differentially expressed in SLGCs after asymmetric division (Ho et al., 2021). Because SPCH is the key player to drive asymmetric cell division and is required for stomatal fate, we further investigated the expression patterns of MYB16-YFP and SPCH-CFP in wild-type (WT) 7 days post-germination (dpg) young true leaves to observe MYB16 behaviors in a stomatal lineage. We checked the asymmetrically divided sister cells, meristemoids and SLGCs, and observed SPCH often localized in meristemoids and MYB16 localized in SLGCs (Figure 1A; Supplemental Figure 1). Further quantification of signals in an entire leaf showed that in a total of 583 pairs analyzed, SPCH was mainly in meristemoids (89.6%), whereas MYB16 was more localized in SLGCs (78.4%) (Figure 1B). However, we also observed MYB16 in a few meristemoids and pavement cells (Supplemental Figure 1). By time-lapse imaging, MYB16 could express in meristemoids, but its expression was quickly replaced by SPCH before asymmetric division (Figure 1C). These observations indicate that MYB16 localization is preferentially in SLGCs, a young cell state in epidermis, with a possible negative relationship between SPCH and MYB16.

**Figure 1.**
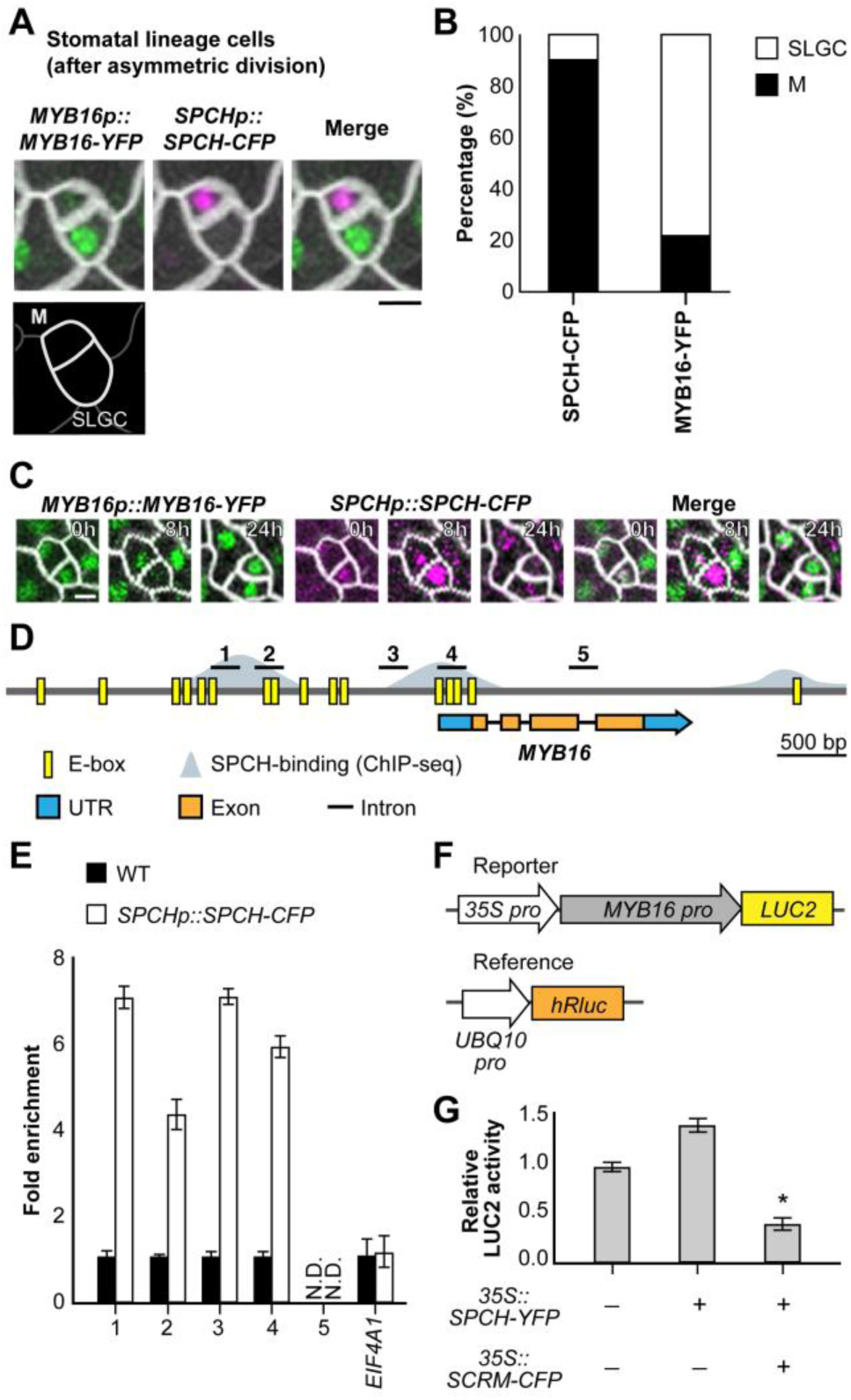
SPEECHLESS (SPCH) binds to *MYB16* promoter and downregulates *MYB16* in meristemoids. **(A)** A still image of MYB16-YFP (green) and SPCH-CFP (magenta) in a meristemoid (M)–stomatal lineage ground cell (SLGC) pair in a 7 days post-germination (dpg) wild-type (WT) true leaf. **(B)** Frequency of SPCH or MYB16 in either cell of meristemoid–SLGC pairs. SPCH is often found in meristemoids as predicted (89.6%). MYB16, in contrast, is preferentially localized to SLGCs (78.4%). A whole leaf image was used to obtain 583 pairs. M, meristemoid; SLGC, stomatal lineage ground cell. **(C)** Time-lapse imaging of SPCH and MYB16 in a meristemoid–SLGC pair. Both SPCH and MYB16 are found in the meristomid at 0 h but only SPCH signal remains at 8 h before asymmetric cell division (24 h). **(D)** Diagram of MYB16 genome region: E-boxes (CANNTG) predicted by PlantPAN 3.0 shown in yellow, SPCH-binding sites obtained from SPCH ChIP-seq data (Lau et al., 2014) shown in gray. Five regions (black bars) designed for the ChIP-qPCR assay were used in **(E)**. **(E)** SPCH binds to the promoter of MYB16 as revealed by ChIP-qPCR assay of 4 dpg *SPCHp::SPCH-CFP* seedlings with GFP-trap beads. Three biological repeats showed similar results. *EIF4A1* is a negative control. N.D., not detected. Data are mean (SD). **(F)** The experimental design for *MYB16* luciferase assay. *MYB16* promoter fused with a mini-35S promoter to enhance the expression. Ratiometric luminescent reporters were used to normalize the expression difference in a given construct. **(G)** SPCH functions with SCRM/ICE1 to downregulate *MYB16* expression. The luciferase assay was performed in 3-week-old WT protoplasts. Four biological repeats showed similar results. *, *p* < 0.001. Data are mean (SD). For **(A)** and **(C)**, cell outlines marked by RC12A-mCherry (gray). Scale bar, 5 µm. See also Supplemental Figure 1.

In line with our hypothesis, SPCH chromatin immunoprecipitation sequencing (ChIP-seq) and induction assays have shown that the promoter of *MYB16* is bound by SPCH and *MYB16* expression is downregulated after SPCH induction (Lau et al., 2014). To confirm the direct binding of SPCH, we searched for the potential SPCH binding motif, E-box, on the *MYB16* promoter by using the PlantPAN 3.0 website-based predictor (Chow et al., 2019) (Figure 1D, yellow boxes). According to the overlap between E-box predictions and the SPCH binding sites derived from Lau et al. (2014) (Figure 1D, grey area), 5 sets of primers were designed for ChIP-qPCR to test SPCH binding to the *MYB16* promoter. Regions 1 to 4 but not the gene body, region 5, showed increased fold change in binding (Figure 1E). To determine whether the binding was positive or negative regulation, we used luciferase assay with ratiometric luminescent reporters to control the expression and copy number of a given construct. A ∼3-kb MYB16 promoter was fused after a mini-35S promoter to enhance the expression (Figure 1F). SPCH is known to form heterodimers with SCRM/ICE1 (Kanaoka et al., 2008), so we included SCRM/ICE1 in the assay as well. SPCH expressed alone in protoplasts conferred no change in luciferase activity as compared with the control. Co-expressing SPCH and SCRM reduced luciferase expression (Figure 1G). In summary, SPCH directly bound to the *MYB16* promoter and suppressed *MYB16* transcription with SCRM/ICE1.

### MYB16 overexpression increased stomatal numbers and clusters

MYB16 functions redundantly with MYB106, so Ho et al. (2021) used a dominant-negative form of MYB16, MYB16-SRDX, to check phenotypes. In contrast to the striking phenotype in organ fusion, MYB16-SRDX showed only slightly decreased stomatal density (stomata/area). In line with the previous observation, the *myb16-crispr* line with a pre-mature stop codon in the first exon also showed slightly reduced stomatal density (Supplemental Figure 2). To further examine whether MYB16 participates in stomatal development, we created several *MYB16* inducible lines. Two, *iMYB16#2 and #3*, showed abnormal stomatal patterns such as clusters and single guard cells under 50 µM β-estradiol treatment for 4 days (Figure 2A to 2H). In contrast to the result from *myb16-crispr*, inducible lines after induction showed higher stomatal density and increased number of stomatal clusters (Figure 2I and 2J). One inducible line, *iMYB16#3*, had more stomata and stomatal clusters under the mock condition (Figure 2F, 2I and 2J) because of the higher MYB16 transcript and protein expression (Figure 2K and 2L). Despite the leaky effect from *iMYB16#3*, overexpression of MYB16 resulted in the formation of stomatal clusters.

**Figure 2.**
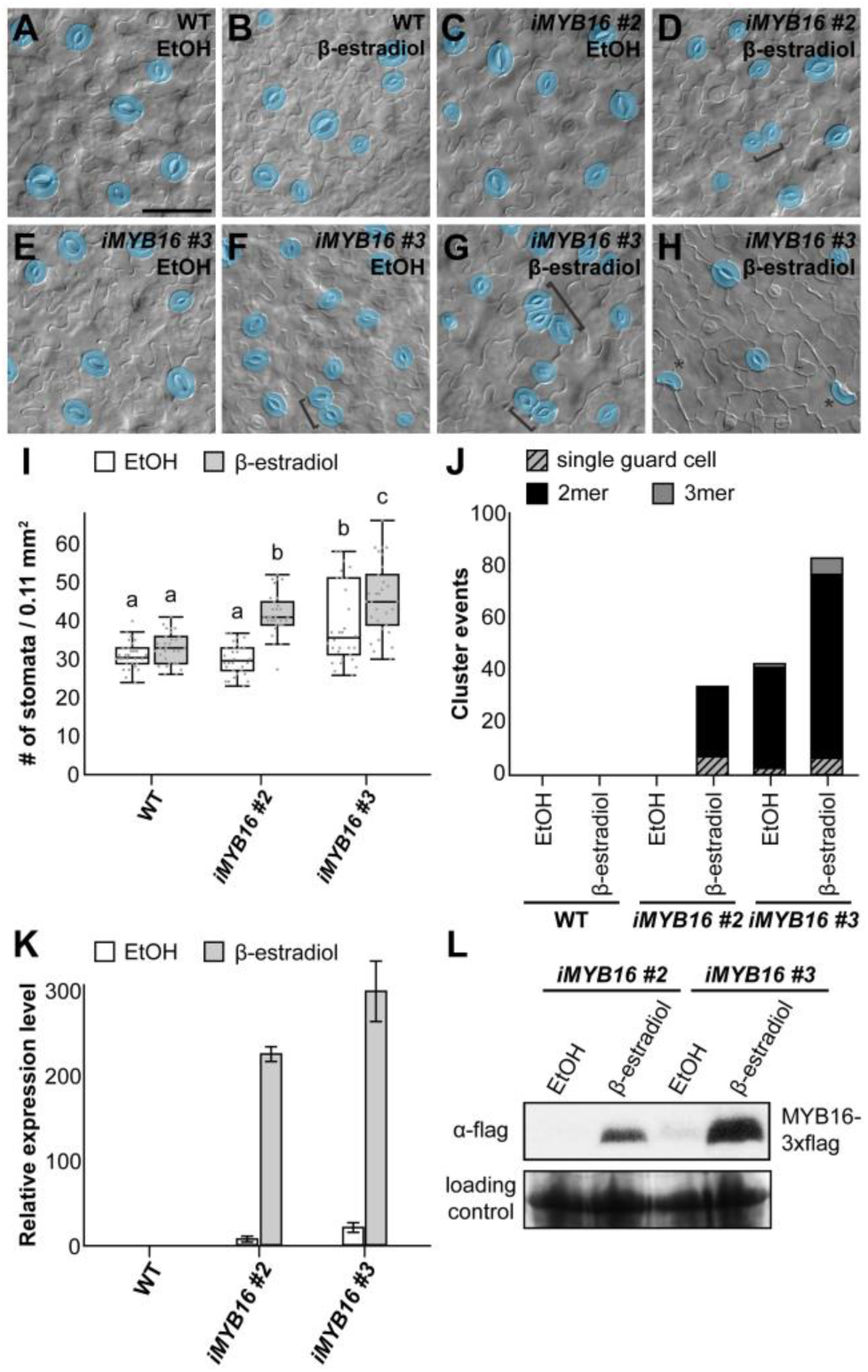
Overexpression of MYB16 induces stomatal clusters. **(A)** to **(H)** DIC images of lower epidermis in 10 dpg WT true leaves and two MYB16 inducible lines (*iMYB16#2* and *#3*) treated with EtOH (mock) or 50 µM β-estradiol. Stomatal pairs (brackets) are often found after β-estradiol treatment in the inducible lines. Occasionally, single guard cells (asterisks) are found **(H)**. Mature guard cells are pseudo-colored in blue. Scale bar, 50 µm. **(I)** Quantification of stomatal density from **(A)** to **(H)**. Stomatal density is increased after β-estradiol treatment in *iMYB16* lines. Compared to *iMYB16#2*, *iMYB16#3* has higher stomatal number under mock treatment, which suggests leaky expression. *p* < 0.05. Kruskal-Wallis test post-hoc with Holm-Bonferroni method. Data are median (interquartile range). **(J)** The quantification of abnormal stomatal phenotypes showing that *iMYB16* lines after β-estradiol treatment have more single guard cell and stomatal clusters (2-3 mer) than WT plants. **(K)** *MYB16* expression detected by qRT-PCR. The expression of MYB16 is induced more than 200 times after β-estradiol treatment. *iMYB16#3* (23.5X) has higher expression then *iMYB16#2* (7.3X) in mock condition. Data are mean (SD). **(L)** Western blot analysis of MYB16 protein level. MYB16 protein level is tightly controlled in *iMYB16#2*. *iMYB16#3* has leaky MYB16 expression in the mock condition, which could explain the phenotypes observed in **(F)**, **(I)** and **(J)**. Coomassie blue staining of total protein is a loading control. A total of 30 lower epidermis samples were observed in **(I)** and **(J)**. See also Supplemental Figure 2.

### Downregulation of MYB16 in meristemoids is required for proper stomatal patterning

To understand the expression timing of MYB16 during stomatal development, we used time-lapse imaging of young true leaves expressing endogenous promoters for MYB16 and SPCH every 8 or 16 h for 4 days for a total of 7 time points. We checked the cell division events and protein localizations in three types of cells — meristemoid, SLGC and protoderm (Figure 3A). The three types of cells showed a similar pattern: co-expression of MYB16 and SPCH or SPCH alone is often found before asymmetric cell division and SPCH remains after the division (Figure 3A). To quantitively analyze the patterns in the time-lapse images, we used the PrefixSpan algorithm (Pei et al., 2001) to explore the possible sequential events of MYB16, SPCH and cell division. A total of 156 cells with strong fluoresce signals were used. Four categories, MYB16-only, SPCH-only, co-localization and division events were noted across the 7 time points (Supplemental Figure 3A). In total, 57 patterns were generated, and the frequency represents how many cells share the pattern (Supplemental Figure 3B). Percentage frequency was calculated by frequency divided by total cell analyzed (frequency/156 cells). First, we focused on the sequential pattern of MYB16 or SPCH expression followed by cell division. The richest pattern was MYB16 and SPCH colocalization before cell division (80.8%) and the next was SPCH alone (70.5%), then MYB16 alone (31.4%) (Figure 3B), which suggests that SPCH but not MYB16 drives cell division. Second, to examine the relation between SPCH and MYB16, we checked all the possible expression combinations and quantified their percentage frequency. The most common phenomenon was MYB16-SPCH colocalization followed by SPCH alone (39.1%) (Figure 3C). Together with our finding of MYB16 inhibition by SPCH, we conclude that SPCH suppresses *MYB16* in meristemoids before asymmetric division. Moreover, although MYB16 expresses in the early state of meristemoids and disappears before asymmetric division, it shows up again in young guard cells (GCs) (Figure 3D and 3G). The tightly regulated expression pattern of MYB16 indicates that the timing of MYB16 downregulation is critical during stomatal development.

**Figure 3.**
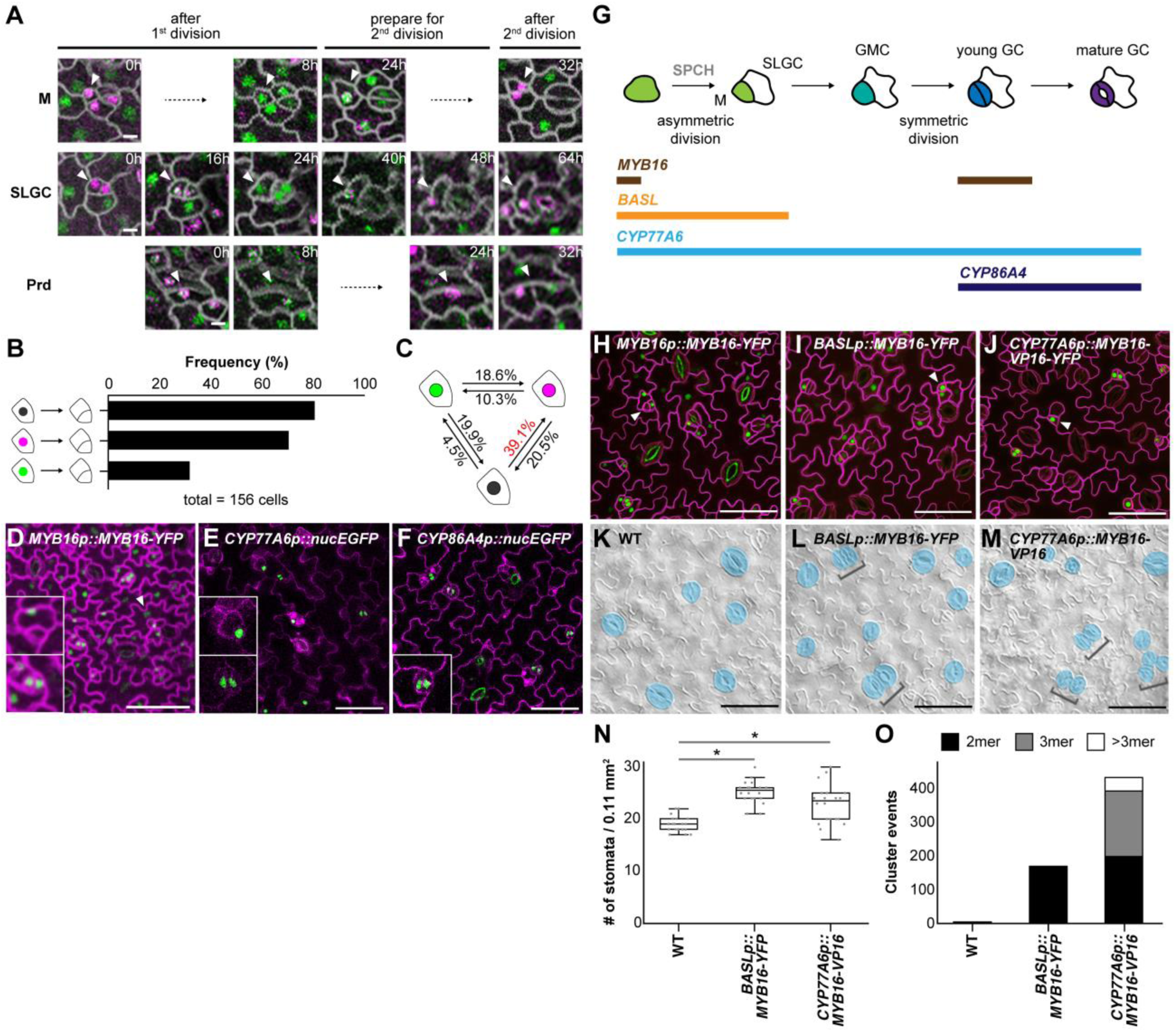
Ectopic expression of MYB16 in early stomatal lineage causes stomatal clusters. **(A)** Time-lapse confocal images of MYB16-GFP (green) and SPCH-CFP (magenta) in 7 dpg true leaves showing that SPCH or MYB16 could express individually or together depending on the sequence of cell division. Arrowheads in different rows indicate cells of interest, in which co-expression of MYB16 and SPCH or SPCH alone is observed before asymmetric cell division and only SPCH remains after the division. Time stamps indicate time since start of the first cell division. M, meristemoid; SLGC, stomatal lineage ground cell; Prd, protoderm. **(B)** Results from the sequential pattern analysis of protein expression using the PrefixSpan algorithm. The colocalization (black) and SPCH alone (magenta) are more frequently seen than is MYB16 alone (green) before cell division. A total of 156 serial events were collected from time-lapse confocal images of 7 dpg true leaves for quantification. **(C)** Frequency of SPCH or MYB16 expression before and after cell division using the PrefixSpan algorithm. The colocalization (black) to SPCH alone (magenta) had the highest frequency (39.1%). **(D)** *MYB16-YFP* driven by the *MYB16* promoter in an 8-dpg true leaf. Confocal images showing MYB16 expression is limited in SLGCs (upper inset), young guard cells (GCs, lower inset) and pavement cells (arrowhead). **(E)** Confocal image of *CYP77A6* transcriptional reporter in 4-dpg cotyledon showing *CYP77A6* expression is stomatal lineage-specific as seen in meristemoid (upper inset) and young guard cells (low inset). **(F)** Confocal image of *CYP86A4* transcriptional reporter in 4-dpg cotyledon showing *CYP86A4* expression is guard cell-specific as seen in young guard cells (inset). **(G)** Summarized expression window of *MYB16*, *BASL*, *CYP77A6* and *CYP86A4* promoters. M, meristemoid; SLGC, stomatal lineage ground cell; GMC, guard mother cell; GC, guard cell. **(H)** to **(J)** Confocal images of 10-dpg true leaves. *MYB16-YFP* is driven by *MYB16* **(H)**, *BASL* **(I)** or *CYP77A6* **(J)** promoter. Compared to SLGC-localized leaves **(H)**, *BASL* and *CYP77A6* promoter-driven *MYB16* **(I)** and **(J)** are seen in meristemoid cells (arrowheads). **(K)** to **(M)** DIC images of lower epidermis from 10 dpg true leaves of WT **(K)**, *BASLp::MYB16-YFP* **(L)** and *CYP77A6p::MYB16-VP16* **(M)** lines. Stomatal clusters (brackets) are found in *BASLp* and *CYP77A6p* lines. Mature guard cells are pseudo-colored in blue. **(N)** Quantification of stomatal density showing increased density when MYB16 is ectopically expressed in stomatal lineage. A total of 20 lower-epidermis samples observed. *, *p* < 0.05, by Wilcoxon signed-rank test. Data are median (interquartile range). **(O)** Total cluster events showing ectopically expressing MYB16 in stomatal lineage causes stomatal clusters. The value obtained from the sum of the events in a total of 20 lower-epidermis samples. Cell outlines marked by *ML1p::RC12A-mCherry* in **(A)**, **(D)** and **(H)** to **(J)** (grey in **[A]**, magenta in **[D]** and **[H]** to **[J]**) and stained by propidium iodide in **(E)** and **(F)**. Scale bars, 5 µm in **(A)** and 50 µm in **(D)** to **(F)** and **(H)** to **(M)**. See also Supplemental Figure 3 to 6.

Stomatal clusters have been found in plants expressing an ectopic form of MYB16, *CYP77A6p::MYB16-VP16.* The form was created by fusion with the promoter from its downstream target, *CYP77A6,* and a transcription activation domain, VP16. Of note, plants expressing MYB16 driven by the promoter of another downstream target, *CYP86A4*, do not show any stomatal phenotype. (Oshima and Mitsuda, 2016) Given that stomatal clusters are in *CYP77A6p::MYB16-VP16* plants only, we wondered whether *CYP77A6* or *CYP86A4* has a distinct expression pattern in epidermis. We analyzed their transcriptional reporters by fusing promoters with nucleus-localized fluorescent protein (nucEGFP) and distinguished stomatal linage cells by staining cell outlines with propidium iodide (PI). Because *CYP77A6* and *CYP86A4* are well-known enzymes in cutin biosynthesis, we expected that they would be expressed in every epidermal cell. However, *CYP77A6* and *CYP86A4* reporters expressed only in stomatal lineage cells (Figure 3E to 3G; Supplemental Figure 4). The fluorescence signal of *CYP77A6p::nucEGFP* was seen in meristemoids, guard mother cells, young GCs and mature GCs with a frequency of 25.6%, 22%, 27.4% and 25%, respectively. The nucEGFP signal of *CYP86A4* was seen only in young and mature GCs, 52.9 and 47.1%, respectively (Supplemental Figure 4). The expression patterns and the stomatal clusters indicate the critical timing of MYB16 expression in the early stage of stomatal development.

BASL expresses before and after asymmetric cell division (Dong et al., 2009); therefore, its promoter as well as the *CYP77A6* promoter would be good choices for ectopic expression of MYB16 in meristemoids (Figure 3G). Similar to previous results, MYB16 expression driven by its native promoter (*MYB16p::MYB16-YFP*) was preferentially localized in SLGCs and young GCs (Figure 3H, SLGC, arrowhead). On ectopically expressing MYB16 by fusion with the *BASL* or *CYP77A6* promoter (*BASLp::MYB16-YFP* or *CYP77A6p::MYB16-VP16-YFP*), nuclear signals were restricted to stomatal lineage cells (Figure 3I and 3J, arrowheads). As compared with WT seedlings, MYB16 ectopic expression lines had higher stomatal density and the increased number of stomatal clusters (Figure 3K to 3O). The high variation of stomatal density observed in *CYP77A6p::MYB16-VP16* was contributed by two types of stomatal phenotypes in seedlings (Supplemental Figure 5). Some of the seedlings had typical clustered stomata and others showed tumor-like colonies (small cell clusters), which is similar to the *basl-1* mutant (Dong et al., 2009). Together with the expression preference and timing of MYB16, the findings suggest that reduction of MYB16 expression in meristemoids is required for proper stomatal formation to prevent stomatal clustering.

### Stomatal clusters in MYB16 ectopic expression lines result from reduced and disrupted localization of polarity protein in asymmetrically dividing cells

Ectopic expression of MYB16 in stomatal lineage results in stomatal clusters. However, we did not observe any significant gene expression changes of stomatal related transcription factors — *SPCH*, *MUTE*, *FAMA* and *SCRM/ICE1* (Supplemental Figure 6). This finding implies the cluster formation caused by signaling cues but not transcriptional regulation. The positional control of polarity complex is required to faithfully place EPF-mediated MAPK signaling in SLGCs to inhibit the stomatal fate (Dong et al., 2009; Houbaert et al., 2018; Pillitteri et al., 2011; Rowe et al., 2019; Zhang et al., 2015)(Figure 4A). To understand how these clusters are formed, we examined cluster formation in plants expressing *BASL* promoter-driven MYB16-YFP and RC12A-mCherry, a cell membrane marker. In the normal condition, a SLGC undergoes asymmetric division to form non-adjacent stomatal precursors (Figure 4B, left). However, frequently seen in the MYB16 ectopic expression line, both sister cells entered symmetric division to form two adjacent stomata (Figure 4B, right).

**Figure 4.**
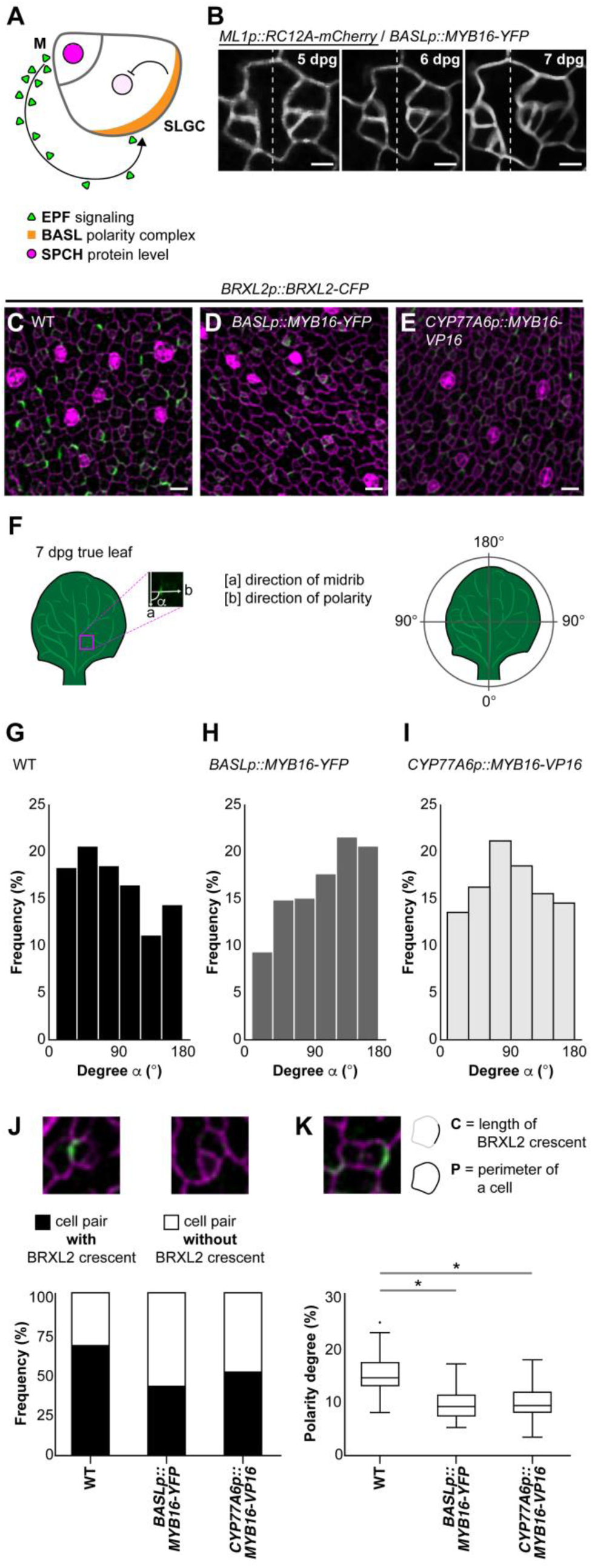
Stomatal clusters are caused by the mis-localization and reduction of polar protein in stomatal lineage. **(A)** Diagram showing the EPIDERMAL PATTERNING FACTOR (EPF)-mediated inhibitory pathway incorporating the spatially labeled polarity complex to prevent stomatal clusters in *Arabidopsis*. EPFs are secreted from meristemoid (M) cells and activate the inhibitory signaling in SLGCs, where the polarity complex recruits inhibitory components, leading to decreased SPCH level. **(B)** Time-lapse confocal images showing stomatal cluster formation in the *BASL* promoter-driven MYB16 ectopic line. Left parts of the images indicate normal stomatal formation. Right parts show the adjacent stoma is derived from an SLGC, resulting in stomatal clustering. **(C)** to **(E)** Confocal images of the polarity marker BRXL2 in 7 dpg true leaves. Compared to WT **(C)**, BRXL2-CFP signal (green) is dimmer in *BASLp::MYB16-YFP* **(D)** and *CYP77A6p::MYB16-VP16* **(E)**. **(F)** Angle α presents the angle between the orientation of midrib and BRXL2. The orientation toward the proximal part of 7 dpg true leaves is set to 0°. **(G)** to **(I)** The orientation of polarity in WT **(G)**, *BASLp::MYB16-YFP* **(H)** and *CYP77A6p::MYB16-VP16* **(I)**. To avoid propidium iodide effect, data were quantified from confocal images of 7 dpg true leaves expressing BRXL2 in the indicated lines without propidium iodide staining. n = 769, 324 and 615 in **(G)** to **(I)**, respectively. **(J)** The presence of BRXL2 in meristemoid–SLGC pairs showing that more cells in WT have BRXL2 crescents. n = 227, 150 and 217 are total cells analyzed in the corresponding order in **(J)**. **(K)** The polarity degree of BRXL2 crescent showing that the WT has the higher polarity degree compared to MYB16 ectopic lines. Polarity degree (C/P) is calculated from crescent length divided by cell perimeter. The dataset is derived from 67 cells with peripheral BRXL2 for each line. The dot shows the Tukey outlier. *, *p* < 0.001, by student t-test. Data are median (interquartile range). Cell outline marked by *ML1p:RC12A-mCherry* in **(B)** and labelled by propidium iodide in **(C)** to **(E)**, **(J)** and **(K)**. Scale bars, 5 µm in **(B)** and 50 µm in **(C)** to **(E)**. See also Supplemental Figure 7.

Disruption of polarity complexes or EPF signaling could cause aberrant stomatal phenotype. We used BRXL2-CFP as an indicator to examine polarity of the polar complex in stomatal cells. The fluorescent intensity of BRXL2-CFP was markedly lower in MYB16 ectopic expression lines than in WT-background plants (Figure 4C to 4E). The upregulated expression of *POLAR*, another factor belonging to the polarity complex, in MYB16 ectopic expression plants might be due to the compensatory effects of polarity loss (Supplemental Figure 6).

To further characterize the polarity defects, we used three parameters to quantify cell polarity. First, we measured the tissue-wise orientation of BRXL2 by the degree α of included angle between the midrib (distal-proximal axis) and cell polarity (Bringmann and Bergmann, 2017) (Figure 4F). The leaves without PI staining were used because we were concerned that the PI solution could damage the cells and further cause polarity change. Most of the BRXL2 orientation was between 30° and 60° in 7-dpg WT seedlings (Figure 4G), as previously observed (Bringmann and Bergmann, 2017). However, in MYB16 ectopic expression lines, BRXL2 orientation peaked between 120° and 150° in *BASLp::MYB16-YFP* and between 60° and 90° in *CYP77A6p::MYB16-VP16* (Figure 4H and 4I). Both MYB16 ectopic expression lines shifted the BRXL2 orientation to the right (larger included angle). Second, we quantified the portion of the cells with or without a BRXL2 crescent in meristemoid–SLGC pairs. The proportion of cells with BRXL2 crescents was decreased in *BASLp::MYB16-YFP* (43%) and *CYP77A6p::MYB16-VP16* (52%) (Figure 4J). Third, the size of the faint BRXL2 signal was smaller in MYB16 ectopic expression lines than in the WT (Figure 4C to 4E). To quantify the crescent size, we used polarity degree (Zhang et al., 2015; Gong et al., 2021) — crescent length normalized by cell perimeter — and found BRXL2 length significantly shorter in both MYB16 ectopic expression lines than in the WT (Figure 4K). In summary, the adjacent stoma in MYB16 ectopic expression lines was derived from an SLGC with reduced and aberrantly oriented BRXL2.

To further confirm that the disrupted polarity establishment causes stomatal clusters in MYB16 ectopic expression lines, we attempted to adjust the polarity defects. Although little is known about how to manipulate polarity degree in plants, one method describes using high-percentage (2.5%) agar to modulate mechanical force (Verger et al., 2018), which may further affect protein polarization. Stomatal density and clusters were rescued under high-percentage agar treatment (Figure 5A and 5B). Moreover, polarity in MYB16 ectopic expression lines was partially rescued (Figure 5C to 5H). With the same quantifying methods used in Figure 4, the orientation as well as degree of polarity were rescued in MYB16 ectopic expression plants grown on high-percentage agar (Figure 5I to 5L).

**Figure 5.**
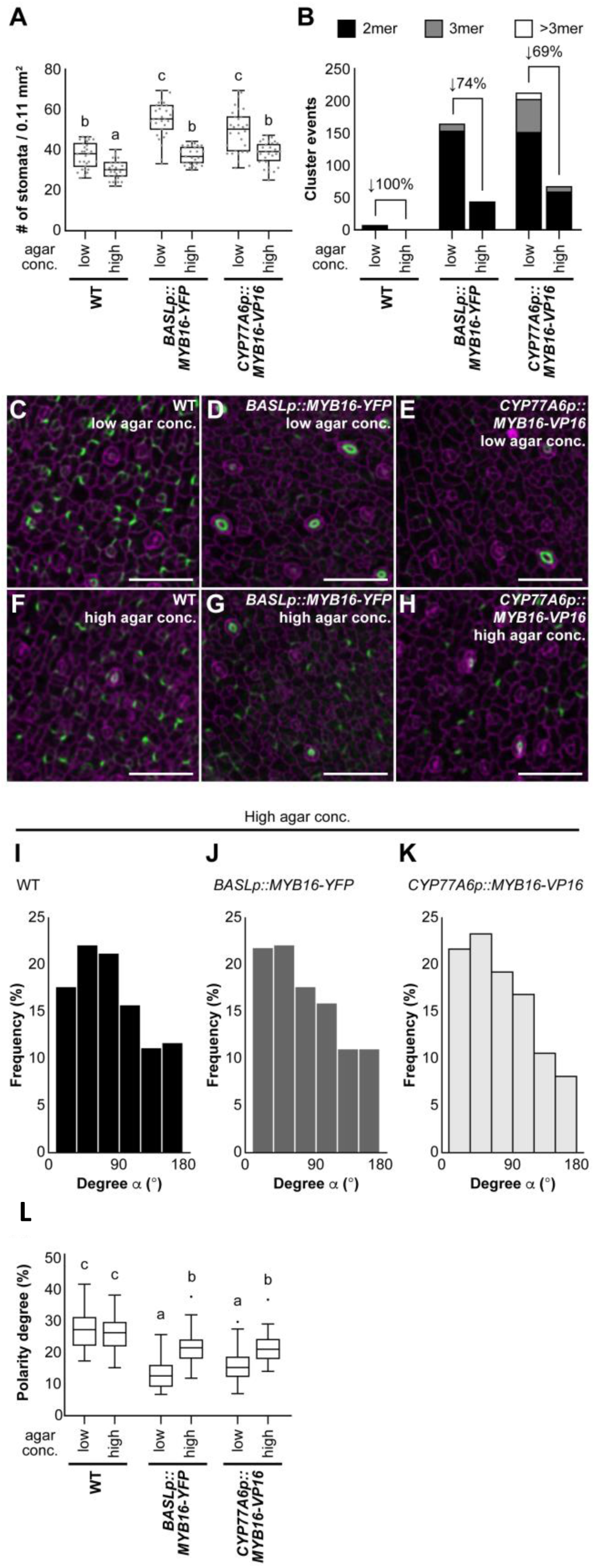
Stomatal phenotype in MYB16 ectopic expression lines was rescued by high-percentage agar treatment. **(A)** Quantification of stomatal density showing the rescue of MYB16 ectopic expression lines by high-percentage agar treatment. Low, 1% agar as normal condition, and high, 2.5% agar. A total of 30 lower-epidermis samples in each 10 dpg plants were observed. Kruskal-Wallis test post-hoc with Holm-Bonferroni method, p < 0.01. Data are median (interquartile range). **(B)** Stomatal clusters are reduced after high-percentage agar treatment. The rescue percentage was calculated by using the difference of cluster events between two kinds of agar treatments divided by the number of cluster events in low-percentage agar treatment. **(C)** to **(H)** Confocal images of the polarity marker BRXL2 in 7-dpg true leaves with two different concentrations of agar treatment. BRXL2-CFP signal (green) is similar in WT **(C)** and **(F)** but stronger in *BASLp::MYB16-YFP* **(D)** and **(G)** and *CYP77A6p::MYB16-VP16* **(E)** and **(H)** after high-percentage agar treatment. **(I)** to **(K)** The orientation of polarity is rescued in *BASLp::MYB16-YFP* and *CYP77A6p::MYB16-VP16* after high-percentage agar treatment. The data were quantified from confocal images of 7-dpg true leaves expressing BRXL2 without propidium iodide staining. n = 587, 369, 321 in **(F)** to **(H)**, respectively. **(L)** The rescue of the BRXL2 crescent size in MYB16 ectopic expression lines by high-percentage agar treatment. The calculated method is the same as in Figure 4K. 60 cells with peripheral BRXL2 of each line under each treatment were collected. The dot shows the Tukey outlier. *p* < 0.001, by two-way ANOVA with Tukey post-hoc test. Data are median (interquartile range). For **(C)** to **(H)**, cell outline is labelled by propidium iodide. Scale bars, 50 µm in **(C)** to **(H)**.

Cell-to-cell signaling is another way to maintain proper distribution of stomata in epidermis. To test whether ectopic expression of MYB16 blocked EPF mediated signaling, we overexpressed EPF2, a major EPF ligand involved in early stomatal development, in *BASLp::MYB16-YFP* plants. Similar to the WT response, the stomatal density was reduced in *BASLp::MYB16-YFP* (Supplemental Figure 7), which suggests that cell-to-cell communication is not affected. Hence, the disruption in polarity in early stomatal development in the MYB16 ectopic expression line was the major reason for clustered stomatal formation.

### Accumulation of cutin affects stomatal development in MYB16 ectopic expression lines

MYB16 is a key regulator of cuticle development (Oshima et al., 2013); therefore, we wondered whether the cuticle was affected in the MYB16 ectopic expression lines. *MYB16* and the cuticular biosynthesis-related genes *LACS1*/*2*, *CYP77A6* and *CYP86A4* were all upregulated in MYB16 ectopic expression lines, with no difference in expression in *GPAT4* and *GPAT8* (Figure 6A and 6B). The cluster phenotypes (Figure 2 and 3) and gene expression of ectopic expression lines were consistent with findings in the MYB16 inducible line (*iMYB16*), with all cuticular biosynthesis-related genes — *LACS1*, *LACS2*, *CYP77A6*, *CYP86A4*, *GPAT4* and *GPAT8* —upregulated under 50 µM β-estradiol treatment (Supplemental Figure 8). The expression of these genes also reflects the degree of stomatal cluster phenotype, with *CYP77A6p::MYB16-VP16* having higher stomatal clusters than *BASLp::MYB16-YFP* (Figure 3O). The correlation of the expression of these enzymes and the amount of cuticle was further supported by toluidine blue (TB) penetration results (Figure 6C). The TB penetration test is often used to measure the permeability of aqueous dye into seedlings (Tanaka et al., 2004). Uptake of TB is easier in seedlings with defective than intact cuticles. The dominant-negative *MYB16p::MYB16-SRDX* plants were strongly stained with TB as compared with WT plants (Figure 6C and 6D). In contrast, the *BASLp::MYB16-YFP* or *CYP77A6p:MYB16-VP16* line had less TB penetration (Figure 6D), which suggests that the cuticular layer is thicker in MYB16 ectopic expression lines.

**Figure 6.**
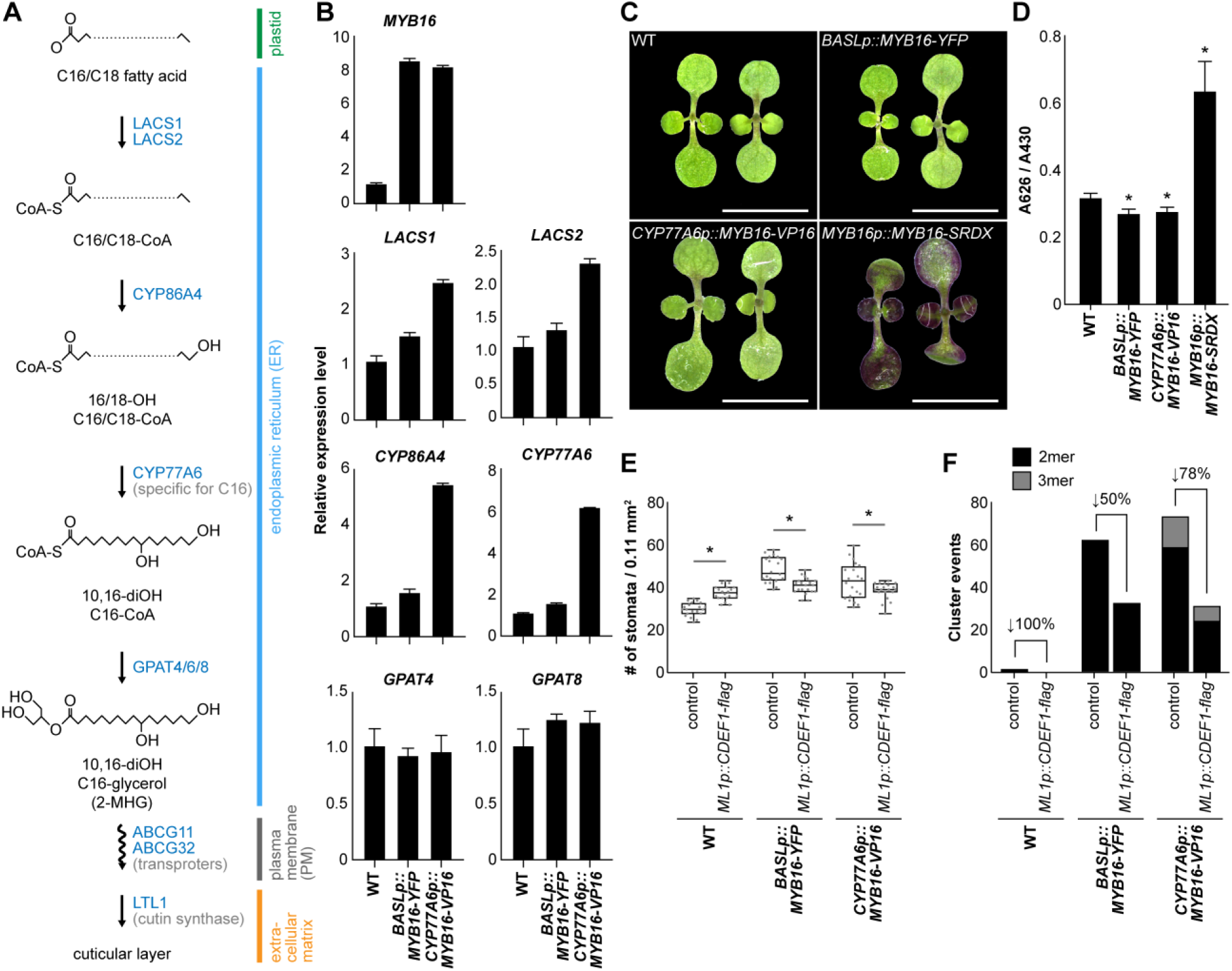
The stomatal clusters in MYB16 ectopic lines are reduced by expression of a cutinase, CDEF1. **(A)** The biosynthesis pathway of the cuticular layer. The C16/C18 fatty acid from plastids is transformed into the acyl-CoA intermediate by LACS1 and LACS2. The hydroxylation of acyl-CoA intermediate is catalyzed by CYP86A4, CYP77A6 and GPAT4/6/8 sequentially to produce the monomer for cuticular layer synthesis. CYP86A4 and GPAT4/6/8 can catalyze both C16 and C18 intermediates, but CYP77A6 preferentially uses C16 as a substrate (Li-Beisson et al., 2009). The transportation of the monomers by ABCG11 and ABCG32 transporters supplies the material required for the polymerization of the cuticular layer outside cell walls. LACS1/2, LONG CHAIN ACYL-COA SYNTHETASE 1/2; GPAT4/6/8, GLYCEROL-3-PHOSPHATE SN-2-ACYLTRANSFERASE 4/6/8. **(B)** Relative mRNA expression of cuticle biosynthesis genes in MYB16 ectopic lines. 7 dpg WT is used as a reference. Data are mean (SD). **(C)** Toluidine blue (TB) test on 7 dpg seedlings of WT, *BASLp::MYB16-YFP*, *CYP77A6p::MYB16-VP16* and *MYB16p::MYB16-SRDX*. Scale bars, 0.5 cm. **(D)** Quantification of penetrated TB showing *MYB16-SRDX* plants are most permeable and *BASLp* or *CYP77A6p* lines are less permeable than WT seedlings. The TB absorbance (A626) is normalized by chlorophyll absorbance (A430). *, *p* < 0.001, by student t-test. **(E)** Quantification of stomatal density showing the partial rescue of MYB16 ectopic expression lines by ectopically expressing cutinase CDEF1. A total of 20 lower epidermis samples in each 10 dpg plants were observed. *, *p* < 0.05, by Wilcoxon signed-rank test. Data are median (interquartile range). **(F)** The stomatal clusters are reduced after ectopically expressing cutinase CDEF1. The rescue percentage is the difference of the cluster event between control and *ML1p::CDEF1-flag* divided by the event number in control. See also Supplemental Figure 8.

To confirm that the formation of stomatal clusters in MYB16 ectopic expression lines is due to the accumulation of cutin, we ectopically expressed CUTICLE DESTRUCTING FACTOR 1 (CDEF1), a cutinase (Takahashi et al., 2010), in epidermal cells by the *ML1* promoter in a *BASLp::MYB16-YFP* or *CYP77A6p:MYB16-VP16* background. The stomatal clusters were reduced in WT plants, and both stomatal density and clusters rescued in MYB16 ectopic expression lines (Figure 6E and F), which suggests the link between stomatal clusters and cuticle production. Stomatal density was also reduced in the *BASLp::MYB16-YFP* or *CYP77A6p:MYB16-VP16* line after introducing CDEF1, but the density was slightly increased in the WT (Figure 6E). These results suggest a crosstalk between stomatal development and cuticle accumulation. MYB16 ectopic expression increased cutin accumulation, which led to abnormal stomatal patterning.

## DISCUSSION

During organ morphogenesis, oriented cell division, extracellular matrix (ECM) and cell–cell signaling are all required for maintaining stem cell multipotency or transforming stem cells to differentiated cells, thereby generating a specific pattern. How the active forces such as signaling cascades and passive forces such as ECM are integrated or regulated in forming a functional tissue remains to be answered. Here, we used an *Arabidopsis* stomatal linage to investigate the coordination of the cuticle layer and fate determiners in leaf epidermal development. By using time-lapse imagining and quantitative analysis of sequential events in lineage progression, we found that MYB16, a key regulator of cutin biosynthesis, expressed in SLGCs, a young state of epidermal cells, and was transcriptionally suppressed in meristemoids by SPCH (Figure 1). One of the functions of an epidermal cell is to produce cuticular layers for a physical barrier, so every protoderm cell in epidermis may be able to express MYB16 to turn on downstream genes for cuticle production. However, the timing of expression is important. We found that for cells in the stomatal lineage, SPCH downregulates MYB16 until the fate is determined. MYB16 appears again in symmetric division in young guard cells whose fate is specified (Figure 3). Eventually a mature stoma with thick cuticular layers is formed. Thus, the formation of cuticle is tightly controlled and integrated with lineage progression to ensure proper development of a stoma.

Many stem cell studies have aimed to understand ECM in determining cell fate. One way is by ECM affecting the stiffness of a tissue to specify the microenvironment and direct stem cell specification (Engler et al., 2006). In animals, mechanobiology has focused on cancer metastasis (Makale, 2007) and tissue regeneration such as hair regeneration in epidermis (Chen et al., 2015). In plants, disruption of ECM reduces cell adhesion and generates disorganized tumor-like growth on shoots (Krupková et al., 2007; San-Bento et al., 2014).Unlike direct cell adhesion mediated by protein linkage and ECM in the animal system, cell adhesion in plants is mostly mediated by the deposition of a pectin-rich middle lamella between adjacent cell walls to promote the mechanical force on epidermis and coordinate the tissue-wise growth. Different from the modification of pectin between cell walls, cuticle acumination occurs mainly on top of leaf epidermis, with possibly covalent linkage between cutin and polysaccharides (Fang et al., 2001). The elevated amount of cuticle may affect overall epidermal mechanical property during leaf expansion. Cuticle production has been linked to stomatal density, but not stomatal clustering yet (Gray et al., 2000; Yang et al., 2011). Because *MYB16-crispr* and overexpression lines showed altered stomatal density (Figure 2; Supplemental Figure 2), we cannot rule out that MYB16 could modulate stomatal lineage divisions as well, although SPCH is still the major gene driving the cell division (Figure 3A to 3C). Stomatal polarity has been linked to mechanical force, with BRXL2 localization in a leaf reflecting the growth and tension direction (Bringmann and Bergmann, 2017). The ectopic deposition of cuticle in meristemoids may affect mechanical stress, thus leading to reduced and redistributed polarity complex (Figure 4) and ultimately a change in cell fate (Figure 2 and 3). A suboptimal water condition of high-percentage agar provides a strategy to manipulate mechanical force (Verger et al., 2018) and modulate polarity localization during stomatal development. The rescue of polarity and stomatal clustering under high-agar concentration suggests the disrupted tensile stress in MYB16 ectopic lines (Figure 5). Expression of a cutinase, CDEF1, in epidermis, rescued the cluster phenotypes in MYB16 ectopic lines (Figure 6). All the lines of evidence suggest that ectopic accumulation of cutin on epidermis may affect mechanical stress, which affects the polarity establishment during stomatal development, thus leading to aberrant stomatal patterning. The downregulation of cutin expression in meristemoids is required for correct polarity establishment during asymmetric cell division in stomatal development (Figure 7).

**Figure 7.**
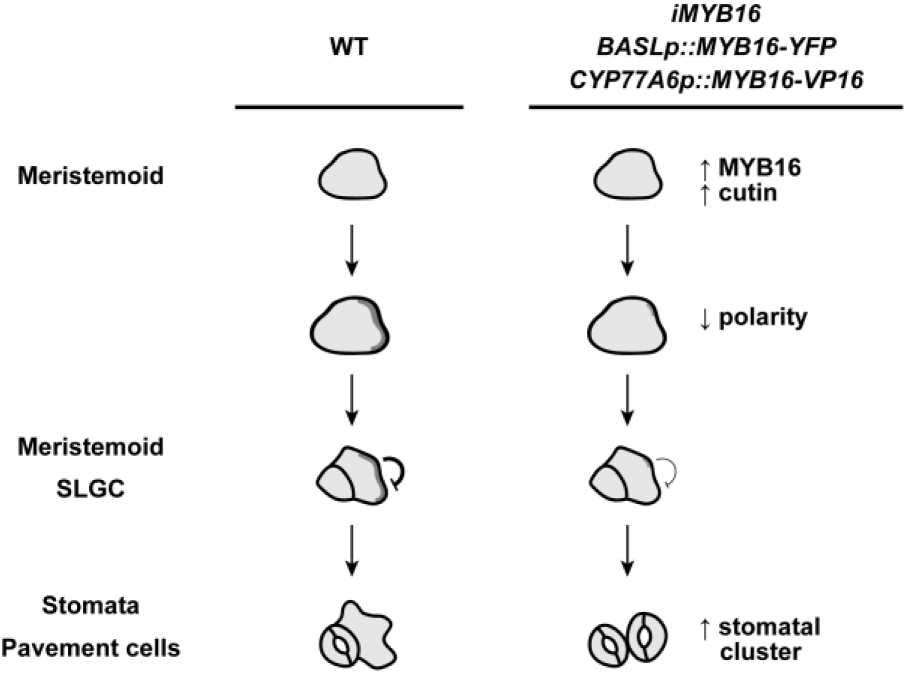
Ectopic MYB16 expression in meristemoids leads to stomatal clusters by modulating polarity protein behavior during asymmetric cell division. In WT epidermis, meristemoids restrict MYB16 expression to ensure polarity establishment for proper stomatal patterning. However, in MYB16 overexpression and ectopic expression lines, high MYB16 expression in meristemoids causes cuticle accumulation. The incorrect timing of cuticle formation may affect the mechanical property in cells, which further causes mis-polarization and reduction of polarity protein during asymmetric cell division. The dark grey shading represents the polarity degree.

Ectopically expressing MYB16 by using lineage-specific promoters allowed for dissecting the critical timing of MYB16 expression (Figure 3G to 3O). Of interest was the stomatal lineage-specific expression patterns of two well-known cutin biosynthesis genes, *CYP77A6* and *CYP86A4* (Figure 3E and 3F; Supplemental Figure 4). CYP77A6 and CYP86A4 are in the cutin biosynthesis pathway according to enzyme functions, mutant phenotypes such as inflorescence fusion and metabolite analysis (Li-Beisson et al., 2009). The mutants *gpat4* and *gpat8* affect stomatal ledge formation (Li et al., 2007), so these two enzymes might be stomatal lineage-specific as well. The finding of stomatal-specific expression patterns of *CYP77A6* and *CYP86A4* suggests that other cutin biosynthesis-related genes might be responsible for cuticle production in pavement cells.

The evolutionary origins of genes that specify stomatal development and function have been resolved phylogenetically in bryophytes (Harris et al., 2020). Genes related to lipid biosynthesis can be traced back to algae. However, the basic cuticle biosynthetic machinery such as CYP77A, GPAT and MYB started to evolve in bryophytes as well (Kong et al., 2020). These lines of evidence together with our findings of *CYP77A6* and *CYP86A4* expression patterns suggest the co-evolution of stomata and cutin machinery in specifying a stoma. One stomatal characteristic is that stomatal ledges, lips around each stomatal pore, are coated with a waterproof layer of cuticle. A mutant, *focl1-1*, showed “fused” ledges and defects in aperture control and transpiration (Hunt et al., 2017). Along with genes involved in structural function, the emergence of MYB transcription factors provides a strategy for spatiotemporal control of gene expression in a developmental context. In *Arabidopsis*, MYB16 and MYB106 regulate cuticle formation in reproductive organs (Oshima et al., 2013). However, MYB16 is a major regulator of cuticle production in vegetative tissues (Oshima and Mitsuda, 2013). The specific function of members in a gene family leads to the diversity of the transcriptional network in building a multicellular organism. With the stomatal linage-specific transcriptomes and single-cell transcriptomes in epidermis (Adrian et al., 2015; Ho et al., 2021; Lopez-Anido et al., 2021), we could start to ask about developmental-regulation and cell type-specific expression of a particular set of genes in cuticle biosynthesis. Biochemical and functional analysis will further pinpoint the metabolic steps in the cuticle synthesis pathway. Modulation of cuticular layers and stomatal numbers on leaf epidermis provides a way to improve plant growth and productivity under drought conditions.

## METHODS

### Plant materials, growth conditions and chemical treatments

Col-0 was the wild type (WT) in all experiments and all transgenic lines were created in this accession. Plant reporters used in this study were *AtML1p::RC12A-mCherry*, *SPCHp::SPCH-CFP* (Davies and Bergmann, 2014), *CYP77A6p::MYB16-VP16*, *MYB16p::MYB16-SRDX* (Oshima et al., 2013). Details for new constructs, *MYB16p::MYB16-YFP, BASLp::MYB16-YFP,* MYB16 inducible system (for *iMYB16*)*, CYP77A6p::MYB16-VP16-YFP, mini35S-MYB16p::LUC2, 35S::SPCH-YFP*, *35S::SCRM-CFP*, *ML1p::CDEF1-flag* and *myb16-crispr* allele are in Supplemental Methods. Primers used for DNA construction are in Supplemental Table 1. Ethanol (EtOH)-sterilized seeds were sown on 1/2 Murashige and Skoog (MS) medium plates and kept at 4°C for stratification. After 24 h, plates were transferred to a 22°C plant tissue culture room with 16 h light/8 h dark cycle. For β-estradiol treatment, β-estradiol in absolute EtOH was added into 1/2 MS medium to the final concentration of 50 µM. An equal volume of absolute EtOH was also added into another 1/2 MS medium as the mock control. Plant materials for phenotyping, mRNA and protein extraction were first grown on 1/2 MS plates for 6 days, then transferred to EtOH or β-estradiol-containing plates for an additional 4 days. Accession numbers are as follows: *MYB16* (AT5G15310), *SPCH* (AT5G53210), *BASL* (AT5G60880), *BRXL2* (AT3G14000), *CYP77A6* (AT3G10570), *CYP86A4* (AT1G01600), *LACS1* (AT2G47240), *LACS2* (AT1G49430), *GPAT4* (AT1G01610), *GPAT8* (AT4G00400) and *CDEF1* (AT4G30140).

### Microscopy

For quantification of stomatal phenotypes, the seedlings were fixed in a 7:1 fixation solution of ethanol and acetic acid for 1 night. Before observation under a Leica DM2500 LED microscopy with DIC prism, the seedlings were washed with MQ water and softened with 1 M KOH until they were completely transparent. The samples were mounted in MQ water for observation. Stomata were quantified from the central region of cotyledons or true leaves. For identifying fluorescence signals, live seedlings were mounted in sterilized water and observed under a Leica STELLARIS 8 (for time-lapse and whole leaf analysis) or Zeiss LSM880 (for expression pattern analysis) confocal microscope. CFP, GFP, and YFP were excited with 458, 488, 514 nm laser, respectively. Propidium iodide (PI) and mCherry were excited with 561 nm laser. PI solution was penetrated into true leaves by applying a vacuum for 30 min.

### mRNA and protein expression

For detection of mRNA expression, total RNA was extracted from plant tissue by using the RNA Plus Mini Kit (LabPrep). Complementary DNA (cDNA) was synthesized from 1 µg total RNA by using SuperScript III transcriptase (Invitrogen). After synthesis, the products were diluted with DEPC-water. The reaction solution contained cDNA template, specific primers and Power SYBR Green Master Mix (Applied Biosystems) for quantitative real-time PCR (qRT-PCR) by using the QuantStudio 12K Flex Real-Time PCR System (Applied Biosystems). The relative expression was analyzed by using QuantStudio 12K Flex software (Applied Biosystems). Primers for qRT-PCR analysis are in Supplemental Table 1.

For protein detection, total protein was extracted from plant tissue by using protein sample buffer (62.5 mM Tris-HCl pH 6.8, 2.5% SDS, 0.002% Bromophenol Blue, 0.7135 M β-mercaptoethanol and 10% glycerol) and boiled under 100°C for 10 min. The extraction was directly used for western blot analysis with 10% SDS-PAGE. After electrophoresis and transferring, the hybridization with proper antibodies by SNAP i.d 2.0 Protein Detection System (Merck) was performed. The chemiluminescence signal was detected in a darkroom by using film. The loading control was the SDS-PAGE stained with Coomassie blue.

### Chromatin immunoprecipitation (ChIP) qRT-PCR assay

To isolate SPCH-chromatin complex, 1.5 g of 4 dpg wild type (Col-0) and *SPCHp::SPCH-CFP* seedlings were harvested and vacuum infiltrated with 1% formaldehyde for 10 min. The crosslinking reaction was then stopped with 2 M glycine solution. After a series steps of homogenizing samples, nuclei isolation, lysis and DNA fragmentation (Haring et al., 2007), the protein–chromatin solution was incubated with GFP antibody-conjugated magnetic beads (ChromoTek) under 4°C overnight. After eluting and reverse crosslinking under 65°C, the final products were purified by using a PCR cleanup column (Geneaid) and eluted in 20 µL 10 mM Tris-HCl (pH 8.0). For qRT-PCR, the reaction solution contained purified products, specific primers and Power SYBR Green Master Mix (Applied Biosystems) mixed and involving the QuantStudio 12K Flex Real-Time PCR System (Applied Biosystems). *EIF4A1* was used as a negative control. The fold enrichment was calculated as follows: Ct value from IP products was divided by Ct value from input, and the ratio was normalized by setting the wild type (Col-0) to 1 on each individual detected region. Primers for qRT PCR analysis are in Supplemental Table 1.

### Luciferase reporter assay

The protoplast transient expression was analyzed by PEG-mediated transformation (Yoo et al., 2007). The reporter, *mini35S-MYB16p::LUC2*, and the effector, *35S::SPCH-YFP* and *35S::SCRM-CFP*, and naked DNA were cotransformed into mesophyll protoplasts isolated from 3-week-old wild-type (Col-0) leaves. After transformation, the protoplasts were incubated in W5 solution under 22°C with one 16 h light/8 h dark cycle. The second day, protoplasts were collected and luciferase activity assay involved the Dual-Luiciferase system (Promega). The relative LUC2 activity was calculated as (luminescence intensity generated by LUC2)/(luminescence intensity generated by Rluc), then values were normalized to the empty vector control, which was set to 1.

### PrefixSpan algorithm

Time-lapse images of 7-dpg seedlings were obtained by using whole-leaf tile scanning with a Leica STELLARIS 8 confocal microscope. The interval time was 8 or 16 h. Cells with serial changes of fluorescence signals and cell division events were recorded (n=156). The matrix was analyzed by using the Python package PrefixSpan, with generator mode (Pei et al., 2001). The output matrix was presented as event counts (Supplemental Figure 2).

### Image processing

For projection of PI-stained confocal images, several stacks were processed by using SurfCut in Fiji-ImageJ (Erguvan et al., 2019).

### Toluidine blue (TB) penetration test

The plants used for the TB penetration test were grown on 1/2 Murashige and Skoog (MS) medium with 0.8% phyto agar (Duchefa Biochemie). An amount of 0.05% (w/v) TB in MQ water was prepared by using a 0.22-µm PVDF filter. Seedlings were immersed with the filtered TB solution for 2 min, then the excessive dye was washed out by using MQ water. Aerial parts were then transferred into tubes containing 1 mL 80% ethanol for 2 h in the dark. The ethanol solution was measured by spectrophotometry with absorbance 430 nm for TB and 626 nm for chlorophyll.

## Supplemental Data

The following materials are available in the online file.

**Supplemental figure 1.** The localization preference of MYB16 is in stomatal lineage ground cells (SLGCs).

**Supplemental figure 2.** Stomatal density is reduced in the *myb16-crispr* mutant.

**Supplemental figure 3.** The workflow of MYB16 and SPCH dynamics by PrefixSpan analysis.

**Supplemental figure 4.** Quantification of CYP77A6 and CYP86A4 transcription patterns.

**Supplemental figure 5.** Two types of stomatal phenotypes in *CYP77A6p::MYB16-VP16*.

**Supplemental figure 6.** Expression of genes related to stomatal development in ectopic MYB16 lines.

**Supplemental figure 7.** Overexpressing EPF2 rescues stomatal clusters in *BASLp*-driven MYB16 lines.

**Supplemental figure 8.** Cuticular-related genes are upregulated after induction in *iMYB16* lines.

**Supplemental Table 1.** Primers used in the study

## ACKNOWLEDGEMENTS

We thank Dr. Yoshimi Oshima and Dr. Nobutaka Mitsuda (National Institute of Advanced Industrial Science and Technology, Japan) for providing *CYP77A6p::MYB16-VP16* and *MYB16p::MYB16-SRDX* lines. We thank Dr. Dominique Bergmann at Stanford University and HHMI, for providing stomata related constructs, Dr. Shu-Hsing Wu at IPMB for providing a 3-FLAG vector, and Dr. Shi-Long Tu and Ping Cheng at IPMB, Academia Sinica, for providing the gateway-compatible destination vector, pCAMBIA1390(GW) containing *35S::XVE-LexA*, for an induction system. We thank Dr. Keng-Hui Lin and Dr. Chih-Wen Yang at Institute of Physics, Academia Sinica and Dr. He-Chun Chou at Research Center for Applied Science, Academia Sinica, for surface tension consulting. We thank Mei-Jane Fang and Ming-Ling Cheng at the Genomic Technology Core Lab (IPMB, Academia Sinica) for DNA sequencing service. We thank Mei-Jane Fang and Ji-Ying Huang at the Cell Biology Core Lab (IPMB, Academia Sinica) for advice on using the Leica STELLARIS 8 and Zeiss LSM880 confocal microscopes. We thank Dr. Paul Verslues (IPMB, Academia Sinica) and Dr. Hsou-Min Li (IMB, Academia Sinica) for their suggestions on this manuscript. This work was supported by the Ministry of Science and Technology in Taiwan (MOST 108-2311-B-001-003-MY3).

## CONTRIBUTIONS OF AUTHORS

S.L.Y. and C.M.K.H. designed experiments. S.L.Y., N.T. and M.Y.T. performed experiments. S.L.Y. and C.M.K.H. wrote the manuscript.

## REFERENCES

Adrian, J. et al. (2015). Transcriptome dynamics of the stomatal lineage: Birth, amplification, and termination of a self-renewing population. Dev. Cell 33: 107–118.

Aharoni, A., Dixit, S., Jetter, R., Thoenes, E., Van Arkel, G., and Pereira, A. (2004). The SHINE clade of AP2 domain transcription factors activates wax biosynthesis, alters cuticle properties, and confers drought tolerance when overexpressed in Arabidopsis. Plant Cell 16: 2463–2480.

Baumann, K., Perez-Rodriguez, M., Bradley, D., Venail, J., Bailey, P., Jin, H., Koes, R., Roberts, K., and Martin, C. (2007). Control of cell and petal morphogenesis by R2R3 MYB transcription factors. Development 134: 1691–1701.

Bessire, M., Borel, S., Fabre, G., Carrac, L., Efremova, N., Yephremov, A., Cao, Y., Jetter, R., Jacquat, A.C., Métraux, J.P., and Nawratha, C. (2011). A member of the PLEIOTROPIC DRUG RESISTANCE family of ATP binding cassette transporters is required for the formation of a functional cuticle in Arabidopsis. Plant Cell 23: 1958–1970.

Bhanot, V., Fadanavis, S.V., and Panwar, J. (2021). Revisiting the architecture, biosynthesis and functional aspects of the plant cuticle: There is more scope. Environ. Exp. Bot. 183: 104364.

Bird, S.M. and Gray, J.E. (2003). Signals from the cuticle affect epidermal cell differentiation. New Phytol. 157: 9–23.

Bringmann, M. and Bergmann, D.C. (2017). Tissue-wide mechanical forces influence the polarity of stomatal stem cells in *Arabidopsis*. Curr. Biol. 27: 877–883.

Broun, P., Poindexter, P., Osborne, E., Jiang, C.Z., and Riechmann, J.L. (2004). WIN1, a transcriptional activator of epidermal wax accumulation in *Arabidopsis*. Proc. Natl. Acad. Sci. USA 101: 4706–4711.

Chen, C.C. et al. (2015). Organ-level quorum sensing directs regeneration in hair stem cell populations. Cell 161: 277–290.

Chow, C.N., Lee, T.Y., Hung, Y.C., Li, G.Z., Tseng, K.C., Liu, Y.H., Kuo, P.L., Zheng, H.Q., and Chang, W.C. (2019). PlantPAN3.0: A new and updated resource for reconstructing transcriptional regulatory networks from ChIP-seq experiments in plants. Nucleic Acids Res. 47: D1155–D1163.

Davies, K.A. and Bergmann, D.C. (2014). Functional specialization of stomatal bHLHs through modification of DNA-binding and phosphoregulation potential. Proc. Natl. Acad. Sci. USA 111: 15585–15590.

Dong, J., MacAlister, C.A., and Bergmann, D.C. (2009). BASL controls asymmetric cell division in Arabidopsis. Cell 137: 1320–1330.

Engler, A.J., Sen, S., Sweeney, H.L., and Discher, D.E. (2006). Matrix elasticity directs stem cell lineage specification. Cell 126: 677–689.

Erguvan, Ö., Louveaux, M., Hamant, O., and Verger, S. (2019). ImageJ SurfCut: A user-friendly pipeline for high-throughput extraction of cell contours from 3D image stacks. BMC Biol. 17: 38.

Fang, X., Qiu, F., Yan, B., Wang, H., Mort, A.J., and Stark, R.E. (2001). NMR studies of molecular structure in fruit cuticle polyesters. Phytochemistry 57: 1035–1042.

Galletti, R., Verger, S., Hamant, O., and Ingram, G.C. (2016). Developing a ‘thick skin’: A paradoxical role for mechanical tension in maintaining epidermal integrity? Development 143: 3249–3258.

Geisler, M., Nadeau, J., and Sack, F.D. (2000). Oriented asymmetric divisions that generate the stomatal spacing pattern in Arabidopsis are disrupted by the *too many mouths* mutation. Plant Cell 12: 2075–2086.

Gong, Y., Varnau, R., Wallner, E.S., Acharya, R., Bergmann, D.C., and Cheung, L.S. (2021). Quantitative and dynamic cell polarity tracking in plant cells. New Phytol. 230: 867–877.

Gray, J.E., Holroyd, G.H., Van Der Lee, F.M., Bahrami, A.R., Sijmons, P.C., Woodward, F.I., Schuch, W., and Hetherington, A.M. (2000). The HIC signalling pathway links CO2 perception to stomatal development. Nature 408: 713–716.

Hara, K., Kajita, R., Torii, K.U., Bergmann, D.C., and Kakimoto, T. (2007). The secretory peptide gene EPF1 enforces the stomatal one-cell-spacing rule. Genes Dev. 21: 1720–1725.

Haring, M., Offermann, S., Danker, T., Horst, I., Peterhansel, C., and Stam, M. (2007). Chromatin immunoprecipitation: Optimization, quantitative analysis and data normalization. Plant Methods 3: 1–16.

Harris, B.J., Harrison, C.J., Hetherington, A.M., and Williams, T.A. (2020). Phylogenomic evidence for the monophyly of Bryophytes and the reductive evolution of stomata. Curr. Biol. 30: 2001–2012.

Heisler, M.G., Hamant, O., Krupinski, P., Uyttewaal, M., Ohno, C., Jönsson, H., Traas, J., and Meyerowitz, E.M. (2010). Alignment between PIN1 polarity and microtubule orientation in the shoot apical meristem reveals a tight coupling between morphogenesis and auxin transport. PLoS Biol. 8: e1000516.

Ho, C.K., Bringmann, M., Oshima, Y., Mitsuda, N., and Bergmann, D.C. (2021). Transcriptional profiling reveals signatures of latent developmental potential in Arabidopsis stomatal lineage ground cells. Proc. Natl. Acad. Sci. USA 118: e2021682118.

Houbaert, A., Zhang, C., Tiwari, M., Wang, K., de Marcos Serrano, A., Savatin, D.V., Urs, M.J., Zhiponova, M.K., Gudesblat, G.E., Vanhoutte, I., et al. (2018). POLAR-guided signalling complex assembly and localization drive asymmetric cell division. Nature 563, 574–578.

Houk, A.R., Jilkine, A., Mejean, C.O., Boltyanskiy, R., Dufresne, E.R., Angenent, S.B., Altschuler, S.J., Wu, L.F., and Weiner, O.D. (2012). Membrane tension maintains cell polarity by confining signals to the leading edge during neutrophil migration. Cell 148: 175–188.

Hunt, L., Amsbury, S., Baillie, A., Movahedi, M., Mitchell, A., Afsharinafar, M., Swarup, K., Denyer, T., Hobbs, J.K., Swarup, R., Fleming, A.J., and Gray, J.E. (2017). Formation of the stomatal outer cuticular ledge requires a guard cell wall proline-rich protein. Plant Physiol. 174: 689–699.

Hunt, L. and Gray, J.E. (2009). The signaling peptide EPF2 controls asymmetric cell divisions during stomatal development. Curr. Biol. 19: 864–869.

Kanaoka, M.M., Pillitteri, L.J., Fujii, H., Yoshida, Y., Bogenschutz, N.L., Takabayashi, J., Zhu, J.K., and Torii, K.U. (2008). SCREAM/ICE1 and SCREAM2 specify three cell-state transitional steps leading to Arabidopsis stomatal differentiation. Plant Cell 20: 1775–1785.

Kong, L., Liu, Y., Zhi, P., Wang, X., Xu, B., Gong, Z., and Chang, C. (2020). Origins and evolution of cuticle biosynthetic machinery in land plants. Plant Physiol. 184: 1998–2010.

Krupková, E., Immerzeel, P., Pauly, M., and Schmülling, T. (2007). The TUMOROUS SHOOT DEVELOPMENT2 gene of Arabidopsis encoding a putative methyltransferase is required for cell adhesion and co-ordinated plant development. Plant J. 50: 735–750.

Lampard, G.R., MacAlister, C.A., and Bergmann, D.C. (2008). Arabidopsis stomatal initiation is controlled by MAPK-mediated regulation of the bHLH SPEECHLESS. Science 322: 1113–1116.

Lau, O.S., Davies, K.A., Chang, J., Adrian, J., Rowe, M.H., Ballenger, C.E., and Bergmann, D.C. (2014). Direct roles of SPEECHLESS in the specification of stomatal self-renewing cells. Science 345: 1605–1609.

Lee, J.S., Kuroha, T., Hnilova, M., Khatayevich, D., Kanaoka, M.M., Mcabee, J.M., Sarikaya, M., Tamerler, C., and Torii, K.U. (2012). Direct interaction of ligand-receptor pairs specifying stomatal patterning. Genes Dev. 26: 126–136.

Li-Beisson, Y., Pollard, M., Sauveplane, V., Pinot, F., Ohlrogge, J., and Beisson, F. (2009). Nanoridges that characterize the surface morphology of flowers require the synthesis of cutin polyester. Proc. Natl. Acad. Sci. USA 106: 22008–22013.

Li, Y., Beisson, F., Koo, A.J.K., Molina, I., Pollard, M., and Ohlrogge, J. (2007). Identification of acyltransferases required for cutin biosynthesis and production of cutin with suberin-like monomers. Proc. Natl. Acad. Sci. USA 104: 18339–18344.

Lopez-Anido, C.B., Vatén, A., Smoot, N.K., Sharma, N., Guo, V., Gong, Y., Anleu Gil, M.X., Weimer, A.K., and Bergmann, D.C. (2021). Single-cell resolution of lineage trajectories in the Arabidopsis stomatal lineage and developing leaf. Dev. Cell 56: 1043–1055.

Lü, S., Song, T., Kosma, D.K., Parsons, E.P., Rowland, O., and Jenks, M.A. (2009). Arabidopsis CER8 encodes LONG-CHAIN ACYL-COA SYNTHETASE 1 (LACS1) that has overlapping functions with LACS2 in plant wax and cutin synthesis. Plant J. 59: 553–564.

MacAlister, C.A., Ohashi-Ito, K., and Bergmann, D.C. (2007). Transcription factor control of asymmetric cell divisions that establish the stomatal lineage. Nature 445: 537–540.

Makale, M. (2007). Cellular mechanobiology and cancer metastasis. Birth Defects Res. 81: 329–343.

Matas, A.J., Cobb, E.D., Bartsch, J.A., Paolillo, D.J., and Niklas, K.J. (2004). Biomechanics and anatomy of *Lycopersicon esculentum* fruit peels and enzyme-treated samples. Am. J. Bot. 91: 352–360.

McFarlane, H.E., Shin, J.J.H., Bird, D.A., and Samuelsa, A.L. (2010). Arabidopsis ABCG transporters, which are required for export of diverse cuticular lipids, dimerize in different combinations. Plant Cell 22: 3066–3075.

Oshima, Y. and Mitsuda, N. (2016). Enhanced cuticle accumulation by employing MIXTA-like transcription factors. Plant Biotechnol. 33: 161–168.

Oshima, Y. and Mitsuda, N. (2013). The MIXTA-like transcription factor MYB16 is a major regulator of cuticle formation in vegetative organs. Plant Signal. Behav. 8: e26826.

Oshima, Y., Shikata, M., Koyama, T., Ohtsubo, N., Mitsuda, N., and Ohme-Takagi, M. (2013). MIXTA-like transcription factors and WAX INDUCER1/SHINE1 coordinately regulate cuticle development in *Arabidopsis* and *Torenia fournieri*. Plant Cell 25: 1609–1624.

Pei, J., Han, J., Mortazavi-Asl, B., Pinto, H., Chen, Q., Dayal, U., and Hsu, M.C. (2001). PrefixSpan: Mining sequential patterns efficiently by prefix-projected pattern growth. Proc. 17th Int. Conf. Data Eng.

Pillitteri, L.J., Peterson, K.M., Horst, R.J., and Torii, K.U. (2011). Molecular profiling of stomatal meristemoids reveals new component of asymmetric cell division and commonalities among stem cell populations in *Arabidopsis*. Plant Cell 23: 3260–3275.

Rowe, M.H., Dong, J., Weimer, A.K., and Bergmann, D.C. (2019). A plant-specific polarity module establishes cell fate asymmetry in the Arabidopsis stomatal lineage. bioRxiv doi: 10.1101/614636.

San-Bento, R., Farcot, E., Galletti, R., Creff, A., and Ingram, G. (2014). Epidermal identity is maintained by cell-cell communication via a universally active feedback loop in *Arabidopsis thaliana*. Plant J. 77: 46–58.

Stracke, R., Werber, M., and Weisshaar, B. (2001). The R2R3-MYB gene family in *Arabidopsis thaliana*. Curr. Opin. Plant Biol. 4: 447–456.

Takahashi, K., Shimada, T., Kondo, M., Tamai, A., Mori, M., Nishimura, M., and Hara-Nishimura, I. (2010). Ectopic expression of an esterase, which is a candidate for the unidentified plant cutinase, causes cuticular defects in *Arabidopsis thaliana*. Plant Cell Physiol. 51: 123–131.

Tanaka, T., Tanaka, H., Machida, C., Watanabe, M., and Machida, Y. (2004). A new method for rapid visualization of defects in leaf cuticle reveals five intrinsic patterns of surface defects in *Arabidopsis*. Plant J. 37: 139–146.

Verger, S., Long, Y., Boudaoud, A., and Hamant, O. (2018). A tension-adhesion feedback loop in plant epidermis. eLife 7: 1–25.

Wang, H., Ngwenyama, N., Liu, Y., Walker, J.C., and Zhang, S. (2007). Stomatal development and patterning are regulated by environmentally responsive mitogen-activated protein kinases in *Arabidopsis*. Plant Cell 19: 63–73.

Yang, J., Isabel Ordiz, M., Jaworski, J.G., and Beachy, R.N. (2011). Induced accumulation of cuticular waxes enhances drought tolerance in *Arabidopsis* by changes in development of stomata. Plant Physiol. Biochem. 49: 1448–1455.

Yang, W., Pollard, M., Li-Beisson, Y., Beisson, F., Feig, M., and Ohlrogge, J. (2010). A distinct type of glycerol-3-phosphate acyltransferase with sn-2 preference and phosphatase activity producing 2-monoacylglycerol. Proc. Natl. Acad. Sci. USA 107: 12040–12045.

Yang, W., Simpson, J.P., Li-Beisson, Y., Beisson, F., Pollard, M., and Ohlrogge, J.B. (2012). A land-plant-specific glycerol-3-phosphate acyltransferase family in Arabidopsis: Substrate specificity, *sn*-2 preference, and evolution. Plant Physiol. 160: 638–652.

Yeats, T.H. and Rose, J.K.C. (2013). The formation and function of plant cuticles. Plant Physiol. 163: 5–20.

Yoo, S.D., Cho, Y.H., and Sheen, J. (2007). Arabidopsis mesophyll protoplasts: A versatile cell system for transient gene expression analysis. Nat. Protoc. 2: 1565–1572.

Zeiger, E. and Stebbins, G.L. (1972). Developmental genetics in barley: a mutant for stomatal development. Am. J. Bot. 59: 143–148.

Zhang, Y., Wang, P., Shao, W., Zhu, J.K., and Dong, J. (2015). The BASL polarity protein controls a MAPK signaling feedback loop in asymmetric cell division. Dev. Cell 33: 136–149.

